# Six-Dimensional Single-Molecule Imaging with Isotropic Resolution using a Multi-View Reflector Microscope

**DOI:** 10.1101/2022.06.26.497661

**Authors:** Oumeng Zhang, Zijian Guo, Yuanyuan He, Tingting Wu, Michael D. Vahey, Matthew D. Lew

## Abstract

We report a radially and azimuthally polarized multi-view reflector (raMVR) microscope for precise imaging of the 3D positions and 3D orientations of single molecules (SMs, 10.9 nm and 2.0° precisions using 5000 photons). These precisions are ∼1.5 times better than those of existing methods for SM orientation-localization microscopy. The raMVR microscope achieves 6D super-resolution imaging of Nile red (NR) molecules transiently bound to 150 nm, 350 nm, and 1 µm-diameter lipid-coated spheres, accurately resolving their spherical morphology despite refractive-index mismatch. Simply by observing the rotational dynamics o raMVR images also resolve the infiltration of lipid membranes by amyloid-beta oligomers without covalent labeling. Finally, we demonstrate 6D imaging of HEK-293T cell membranes, where the orientations of merocyanine 540 molecules reveal heterogeneities in membrane fluidity. With its ∼2 µm depth range, nearly isotropic 3D spatial resolution, and superior orientation measurement precision, we expect the raMVR microscope to enable 6D imaging of molecular dynamics within biological and chemical systems with unprecedented detail.

## Introduction

A hallmark of biological systems is their careful control and regulation of biomolecular interactions at the nanoscale, via mechanical forces, ^1^ lipid membranes, ^2^ and biomolecular condensates, ^3,4^ to name a few examples. Owing to their sensitivity and versatility, ^5,6^ single fluorescent molecules and their translational and rotational dynamics have been used to study a variety of biochemical processes, including the motions of molecular motors, ^7,8^ conformations of supercoiled DNA, ^9–11^ and the structure and organization of actin filaments, ^12,13^ amyloid aggregates, ^14,15^ and lipid membranes. ^16,17^ Simultaneously resolving the 3D position, 3D orientation, and rotational diffusion of single fluorophores, termed single molecule orientation localization microscopy (SMOLM), requires careful manipulation of the phase and polarization of only hundreds or thousands of photons. Limited by challenging signal-to-background ratios (SBRs) in typical single molecule (SM) experiments, microscopists have developed various SMOLM techniques to improve measurement performance, ^18^ approaching the theoretical limits under ideal conditions. ^19–21^ To use precious fluorescence photons more efficiently and maximize signal-to-noise ratios (SNRs), most recently developed techniques create dipole-spread functions (DSFs), or images of dipole emitters, with compact footprints ^11,12,15,17,22^ and thusly achieve higher precision and accuracy. One key challenge with these designs is that orientation estimation requires fitting subtle features within noisy images. Therefore, optical aberrations must be carefully calibrated, often requiring online measurement of specific sample-induced aberrations. ^23,24^ In contrast, orientation-sensitive DSFs based on pupil splitting ^25–27^ are generally more robust against aberrations because their orientation information is encoded in the relative brightness of well-separated “spots.” However, this robustness comes at the cost of typically lower SBRs and worse estimation performance. ^28^

Here, we present a combined polarization modulation and pupil-splitting technique, termed the radially and azimuthally polarized multi-view reflector (raMVR) microscope, that achieves isotropic imaging resolution and ∼1.5 × better precision for measuring 6D SM positions and orientations compared to state-of-the-art techniques while showing excellent robustness against aberrations. We use the raMVR microscope to study model lipid membranes and how amyloid-beta aggregates intercalate within and perturb their structures. We also visualize SM rotational dynamics to reveal heterogeneities in the fluidity of cellular membranes. These data establish the power of raMVR for resolving both the structure and organization of biological targets accurately and precisely with nanoscale resolution.

## Results

### Operating principle and theoretical performance

Fluorescent molecules are well-approximated as oscillating dipole emitters ^29^ (Figure 1b), and we model their rotational diffusion as uniform within a hard-edged cone using an orientation vector ***µ*** = [*µ*_*x*_, *µ*_*y*_, *µ*_*z*_] = [sin *θ* cos *ϕ*, sin *θ* sin *ϕ*, cos *θ*] and solid angle Ω ∈ [0, 2*π*]. To measure molecular orienta-tion, our raMVR approach takes inspiration from both polarization modulation ^12,17,30^ and pupil-splitting ^25,26^ innovations in SM orientation-localization microscopy (SMOLM). First, the polarized radiation pattern of a dipole emitter may be manipulated to measure its orientation; ^31–33^ here, we use a vortex (half) waveplate and polarizing beamsplitter (PBS) (Figures 1a) to separate light from emitters oriented parallel to versus perpendicular to the optical axis. ^17,30,34^ Secondly, we wish to maximize orientation measurement sensitivity while simultaneously maintaining superior SBRs in SM imaging. We therefore split fluorescence photons at the back focal plane (BFP) or pupil of the microscope using a square pyramidal mirror (Figure 1c), called the glass pyramid. The resulting four reflected “beams” are directed toward a mirror with four reflective surfaces whose shape is a pyramid hollowed out of a square prism, termed the air pyramid. The apex angle of the air pyramid is slightly smaller compared to that of the glass pyramid (by 1 degree in our system, see Supplementary Section 1) and reflects the fluorescence back in its original direction to create four imaging channels on a camera (Figure 1d). Critically, both signal and background photons are separated across the four channels, thereby maintaining an effective SBR that is similar to a standard unpolarized epifluorescence microscope. While the mirror configuration is similar to that of a Cassegrain reflector (Figure 1c), here we mount our piecewise-planar pyramid mirror from the diagonal directions to avoid blocking any fluorescence (see Figure S1), achieving a photon efficiency close to 100% and superior to that of metasurfaces ^30^ and spatial light modulators. ^15,25–27^

**Figure 1:**
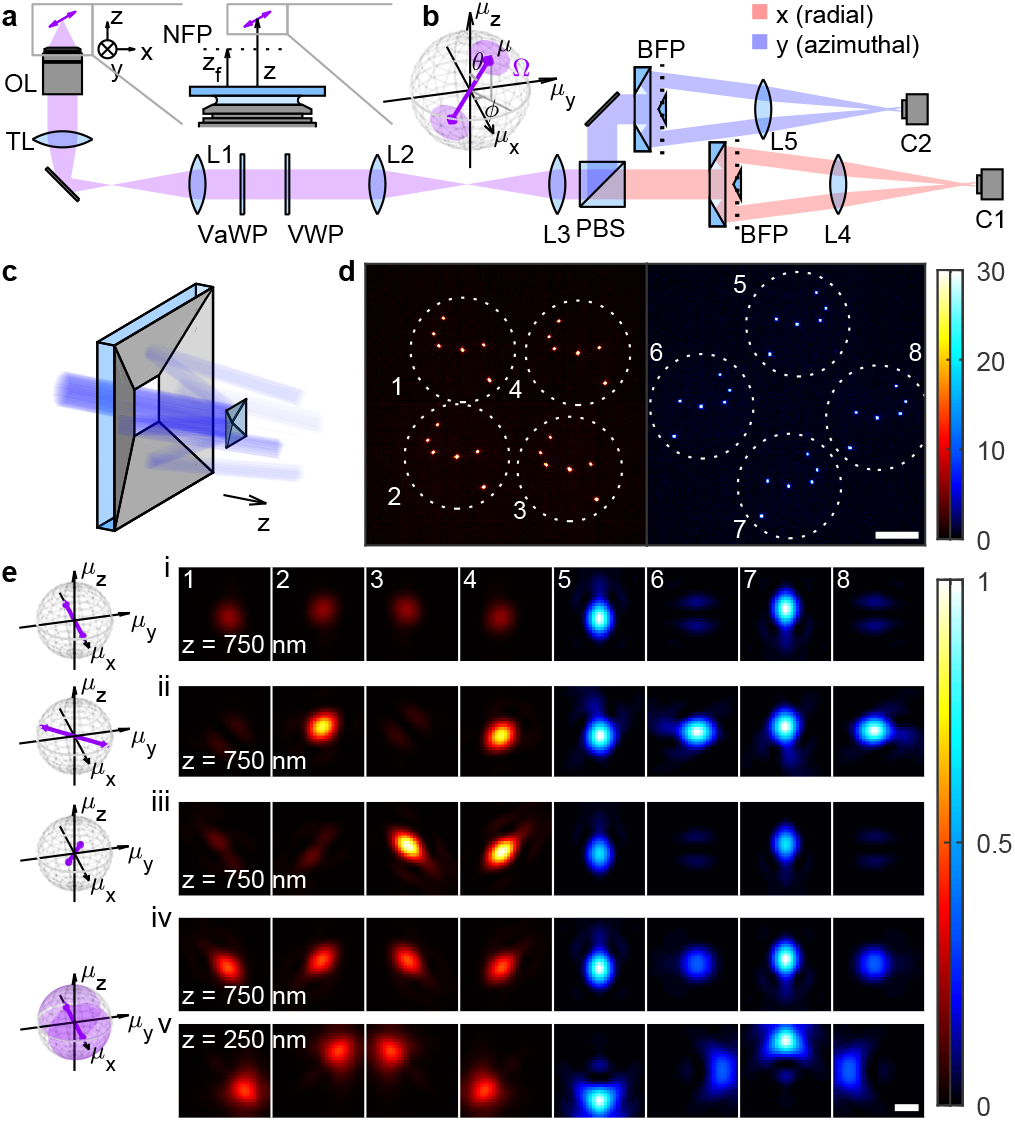
Concept of raMVR single-molecule orientation-localization microscopy (SMOLM). (a) Schematic of the radially and azimuthally polarized multi-view reflector (raMVR) microscope. (b) The fluorescent molecule is modeled as an oscillating dipole uniformly wobbling within a hard-edged cone with an average orientation ***µ*** and a “wobble” solid angle Ω. After turning radially and azimuthally polarized fluorescence to (red) *x*-and (blue) *y*-polarized light using a variable waveplate (VaWP) and a vortex (half) waveplate (VWP), the emission light is separated into different polarization channels using a polarizing beamsplitter (PBS). The (c) raMVR pyramidal mirrors further separate light into four detection channels and project the (d) image plane onto a detector (C1, C2) in each channel. Colorbar: photons; scalebar: 20 µm. (e) Representative DSF in each channel for molecules located at (i-iv) *z* = 750 nm and (v) *z* = 250 nm with orientations (*θ, ϕ*, Ω) of (i) (90°, 0°, 0 sr), (ii) (90°, 45°, 0 sr), (iii) (45°, 0°, 0 sr), and (iv,v) (90°, 0°, *π* sr). Colorbar: normalized intensity; scale-bar: 500 nm; nominal focal plane position *z*_*f*_ = 1200 nm.

Figure 1e shows raMVR images captured from SMs with various orientations. Each SM orientation and degree of rotational diffusion is clearly distinguishable by the varying brightnesses of the focused DSF in each channel. Moreover, because each imaging channel now samples one fourth of the objective’s total aperture, the DSF in each channel *shifts laterally* as the SM’s axial position changes [Figure 1e(iv,v)]. This unique method of encoding both position and orientation information enables the raMVR microscope to achieve estimation precisions [calculated via Cramér-Rao bound (CRB), ^35^ see Supplementary Section 2.3] far superior to that of state-of-the-art methods across a 1.5-µm depth range (Figure 2). With a typical SM SBR of 5000 signal photons (for an in-plane SM at the water-glass interface, *z* = 0) and 40 background photons per 66.8 × 66.8 nm^2^ pixel collected across all imaging channels of the raMVR microscope, the average precision for estimating the average orientation ***µ***, wobble Ω, lateral *x*, and axial *z* positions are 2.04°, 0.23 sr, 10.79 nm, and 12.24 nm, respectively; these numbers are 47.9%, 43.8%, 25.9%, and 55.2% smaller compared to those of CHIDO ^12^ and 66.4%, 65.6%, 42.8%, and 62.0% smaller than those of the vortex DSF. ^11^ Importantly, raMVR’s precision is superior to those of all other orientation-localization methods (see Figure S10 and Table S1 for other comparisons), over its entire depth range, regardless of the SM’s position and orientation. Due to the reduced effective NA with each imaging channel, raMVR exhibits somewhat reduced lateral localization precision; its localization precision is 2.97 nm and 1.23 nm worse than those of CHIDO and the vortex DSF, respectively. However, because of its compact DSF, dim SMs are easier to detect using the raMVR microscope than when using CHIDO and the vortex DSF for defocused SMs and even for certain in-focus SMs (Supplementary Section 2.2). Notably, our detection and estimation algorithm (Supplementary Section 3) maintains these performance advantages on simulated images, attaining an average localization precision of 24% and orientational precision of 5% greater than the CRB (Table S2).

**Figure 2:**
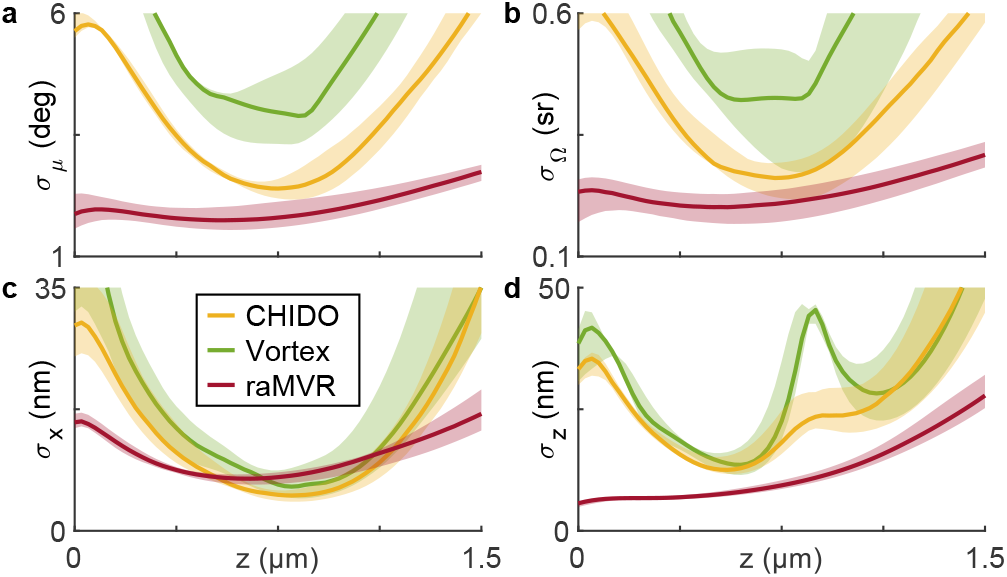
Theoretical precision of measuring the (a) average orientation, (b) wobble angle, (c) lateral position, and (d) axial position of an SM using the raMVR microscope compared to other methods, with 5000 signal photons and 40 background photons per pixel total across all channels. Lines represent the median, and shaded areas represent the 10th and 90th percentiles across all possible orientations. Yellow: CHIDO; green: vortex DSF; red: raMVR.

### Experimental validation of 6D precision and accuracy

Silica spheres coated with supported lipid bilayers (SLBs) composed of DPPC (1,2-Dipalmitoyl-sn-glycero3-phosphocholine) and 40% cholesterol (Figure 3a) have been previously shown ^22^ to be excellent calibration targets for SMOLM since we are able to systematically control both the morphology and ordering of the lipids within the SLB. By imaging Nile red (NR, Figure S15) molecules transiently bound within the SLBs (using the PAINT blinking mechanism ^36^), we resolve spherical structures with radii as small as 150 nm and as large as 1 µm without scanning the sample (Figures 3b,c). Moreover, we observe NR to be perpendicular to the spheres’ surfaces, regardless of the spheres’ radii (Figure 3d), thereby matching previous observations; ^15,16^ the angles *θ*_⊥_ between NR’s average orientation ***µ*** and the sphere’s surface normal vector ***r*** are 12.0° ±15.0°, 13.5°± 17.6°, and 19.0°± 20.4° (median ± std. dev.) for 1000-nm, 350-nm, and 150-nm radius spheres, respectively. We also find that the NR are moderately constrained in their rotational diffusion with wobble cone angles Ω are 1.12 ± 0.93 sr, 1.13 ± 0.95 sr, and 1.01 ± 1.01 sr for 1000-nm, 350-nm, and 150-nm spheres, respectively. We note that despite the severe refractive index mismatch between water (*n*_2_ = 1.33) and the silica sphere supporting each bilayer (*n*_*s*_ = 1.45, Figure 3a), the raMVR system can image NR throughout the entire spherical SLB without any shape distortions (Figure 3b,c), which has never before been achieved by other SMOLM techniques.

**Figure 3:**
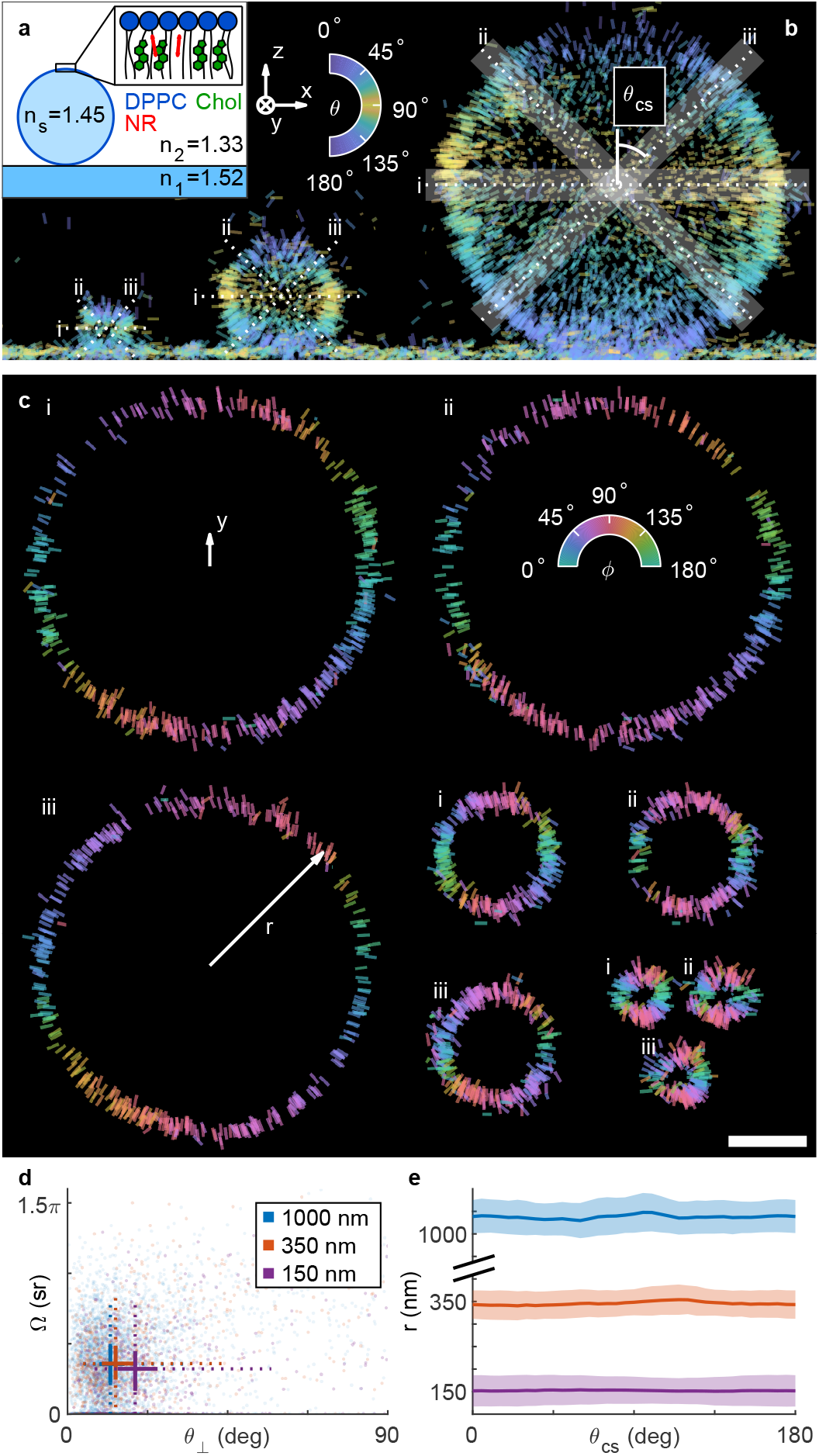
6D nanoscopic imaging of spherical supported lipid bilayers (SLBs). (a) Nile red (NR) molecules transiently binding to DPPC SLBs with 40% cholesterol surrounding a silica sphere. (b) An *xz*-view of all localizations on spheres with radii of 150 nm, 350 nm, and 1000 nm. Lines are oriented and color-coded according to the measured polar angle *θ*. Their lengths are proportional to 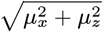 Local-izations within cross sections with tilt angles *θ*_CS_ of (i) 90°, (ii)− 45° and (iii) 45° and thickness of 200 nm. Lines are oriented and color-coded according to the measured azimuthal angle *ϕ*. Their lengths are proportional to the length of the unit vector ***µ*** projected into the cross-sectional planes. (d) NR orientation *θ*_⊥_ = cos^−1^(***µ · r****/r*) and wobble (Ω) measurements for the three spheres in (a). (e) Measured distances *r* from NR localizations to the center of the sphere as a function of the cross-section angle *θ*_CS_. Solid lines represent the average distance, and shaded areas represent ± 1 std. dev. Blue: 1000-nm spheres; orange: 350-nm spheres; purple: 150-nm spheres. Scalebar: 500 nm.

To quantify localization precision, we isolate cross sections of each sphere along various directions *θ*_CS_ (Figure 3b and Movie S1) and measure the distance *r* between the each molecule and the center of the sphere (Figure 3c,e). Because of raMVR’s isotropic localization precision (Figure 2c,d), our measured localization precision is uniform across all cross-section angles *θ*_CS_. The apparent full width at half maximum (FWHM) thicknesses of the SLBs are 76.1 nm for 1000-nm spheres (Movie S1), 42.6 nm for 350-nm spheres, and 51.6 nm for 150-nm spheres (Figure S17). The average distances *r* as functions of *θ*_CS_ yield eccentricities of 0.23, 0.28, and 0.20 for the 1000-nm, 350-nm, and 150-nm spheres, respectively (Figure 3e), where the length ratio between short and long axes is 0.96 for an eccentricity of 0.28. Thus, the raMVR microscope accurately reconstructs spherical structures with nanoscale resolution.

### 6D imaging of interactions between amyloid aggregates and lipid membranes

The nanoscale interactions between lipid membranes and amyloid aggregates are of tremendous interest for studying the pathogenesis of neurodegerative disease, ^37,38^ and while atomic force and transmission electron microscopes have provided detailed static snapshots of these structures, ^39,40^ a 3D imaging technique to resolve the morphology and organization of *individual* peptide-lipid structures is still missing. Here, we use the raMVR system to image aggregates of the 42-residue amyloid-beta peptide (Aβ42), which are a primary signature of Alzheimer’s disease, ^41^ intercalated within spherical SLBs (Figure 4). We incubated Aβ42 monomers with lipid-coated 350-nm spheres at 37°C with constant shaking (200 rpm). While NR binds to both lipid membranes and Aβ42 aggregates, the sensitivity of NR’s rotational dynamics to its chemical environment ^15,16^ enables us to resolve pure SLBs [Figure 4c(i)] from those that have been infiltrated by Aβ42. When in contact with amyloid aggregates on the spheres’ surfaces, NR shows larger wobble angles [Figure 4c(ii-iv), Ω = 2.98 ± 1.14 sr in panel (ii), 2.33 ± 1.20 sr in panel (iii), and 2.09 ± 1.15 sr in panel (iv)] compared to those within DPPC SLBs (Figure 3d, Ω = 1.13 ± 0.95 sr). This larger wobble with Aβ42 present is also easily distinguished from that within a 100-nm pure lipid vesicle attached to a sphere [Figure 4b(i), Ω = 1.69 ± 0.93 sr, and Movie S2]. Kolmogorov-Smirnov tests of these wobble distributions return *p*-values much less than 0.01 (see Supplementary Section 5.2 for details), thereby confirming that NR exhibits dramatically different orientational dynamics when in contact with amyloid-containing SLBs versus pure lipid membranes.

**Figure 4:**
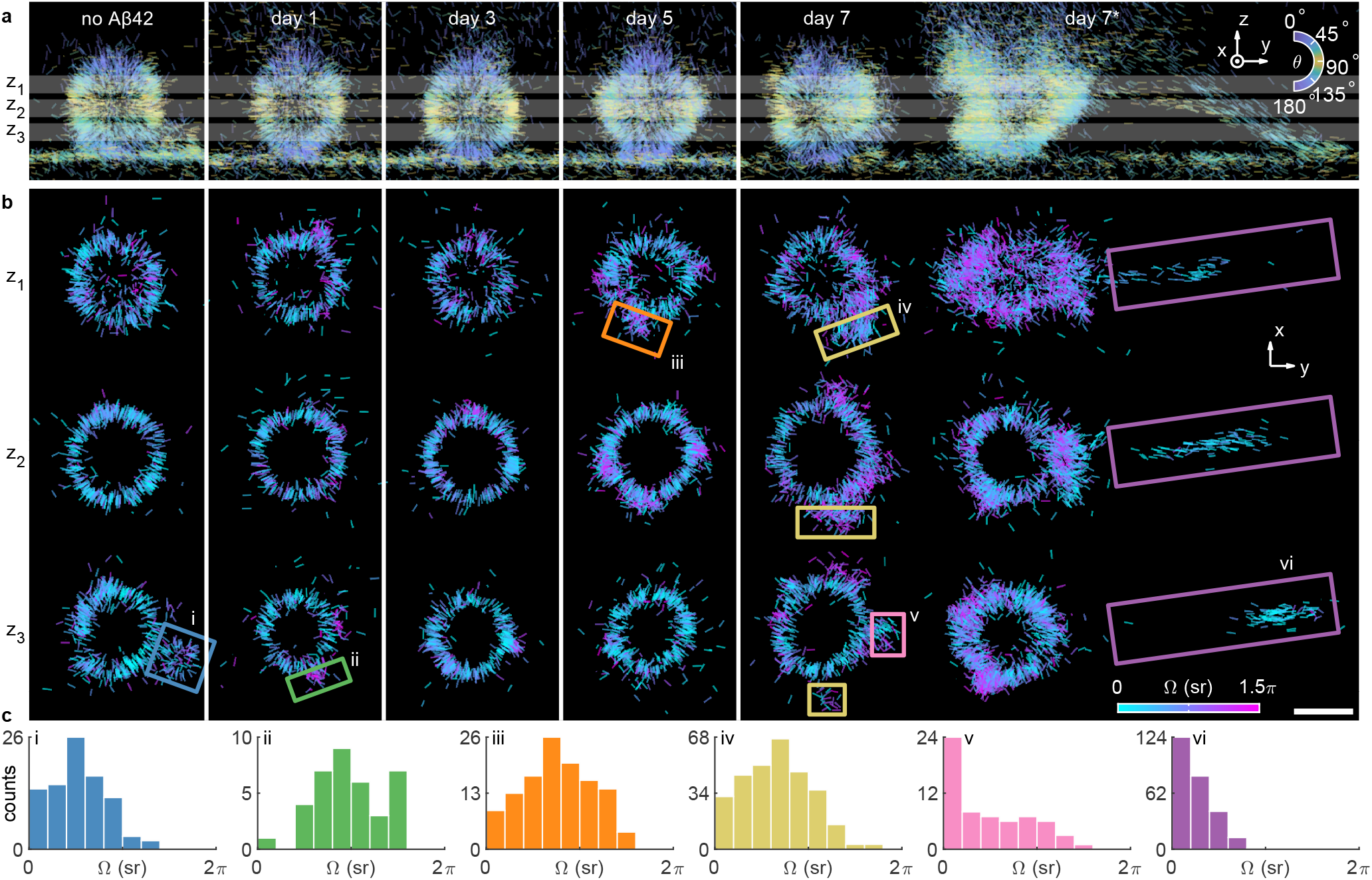
6D nanoscopic imaging of amyloid-lipid interaction. (a) *xz*-views of all NR localizations on 350-nm lipid-coated spheres without (incubated for 7 days) and with (incubated for 1, 3, 5, and 7 days) Aβ42. Lines are oriented and color-coded according to the measured polar angle *θ*. Their lengths are proportional to 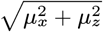. (b) Localizations within the 150 nm-thick *z* slices marked in (a) centered at (*z*_1_ = 600, *z*_2_ = 400, *z*_3_ = 200) nm. Lines are oriented according to the measured azimuthal angle *ϕ* and color-coded according to the measured wobble cone angle Ω. Their lengths are proportional to sin *θ*. Scalebar: 500 nm. (c) Distribution of measured wobble cone angles within the boxed areas in (b).

In the presence of lipids, we observe that Aβ42 is more likely to form oligomers and proto-fibrils than long fibril structures, in agreement with previous studies. ^42,43^ For example, a cluster of NR exhibiting small wobble angles [Figure 4c(v)] reveals a small Aβ42 aggregate attached to the sphere near the coverslip in Figure 4b(v). We also observed larger aggregates linking multiple spheres (Figure S22). Across more than 25 spheres from 7 separate aggregation reactions (Figure S20), we observe only one occurrence of significant fibril growth (Figure 4 and Movie S2, day 7*). NR molecules bound to the fibril appear to be parallel to its long axis and more rotationally-fixed [Figure 4c(vi), Ω = 0.70 ± 0.67 sr] compared to those within the SLBs. This result is consistent with previous work ^14,15^ and a control experiment of Aβ42 fibrils aggregated in the absence of lipids (Figure S14). Overall, principal component analysis of NR orientation distributions (Supplementary Section 5.3) indicates that Aβ42 aggregation in the presence of lipid membranes is extremely heterogeneous (Figures S19-S21); however, membrane disorder generally increases with longer Aβ42 aggregation reactions.

### 6D imaging of the fluidity of HEK-293T cell membranes

Environmentally sensitive fluorophores have exciting promise as molecule-scale sensors for visualizing the nanoscale organization of lipid membranes. ^16,44–48^ We demonstrate, for the first time, 6D imaging of cell membrane morphology and fluidity by analyzing fluorescence flashes of merocyanine 540 (MC540, Figure S15) transiently binding to fixed HEK-293T cells (Figure 5, Movie S3 and Supplementary Section 4). Here, the raMVR microscope measures 3D SM positions and 3D rotational dynamics within a 23 × 23 × 2 µm^3^ volume without scanning the sample or focal plane, obtaining an experimental 3D resolution (using Fourier shell correlation ^49,50^) of 64.1 nm (FWHM) for a vesicle (Figure 5c, yellow box and Figure S23a) and 73.0 nm for a planar membrane (Figure 5c, green box and Figure S23b).

**Figure 5:**
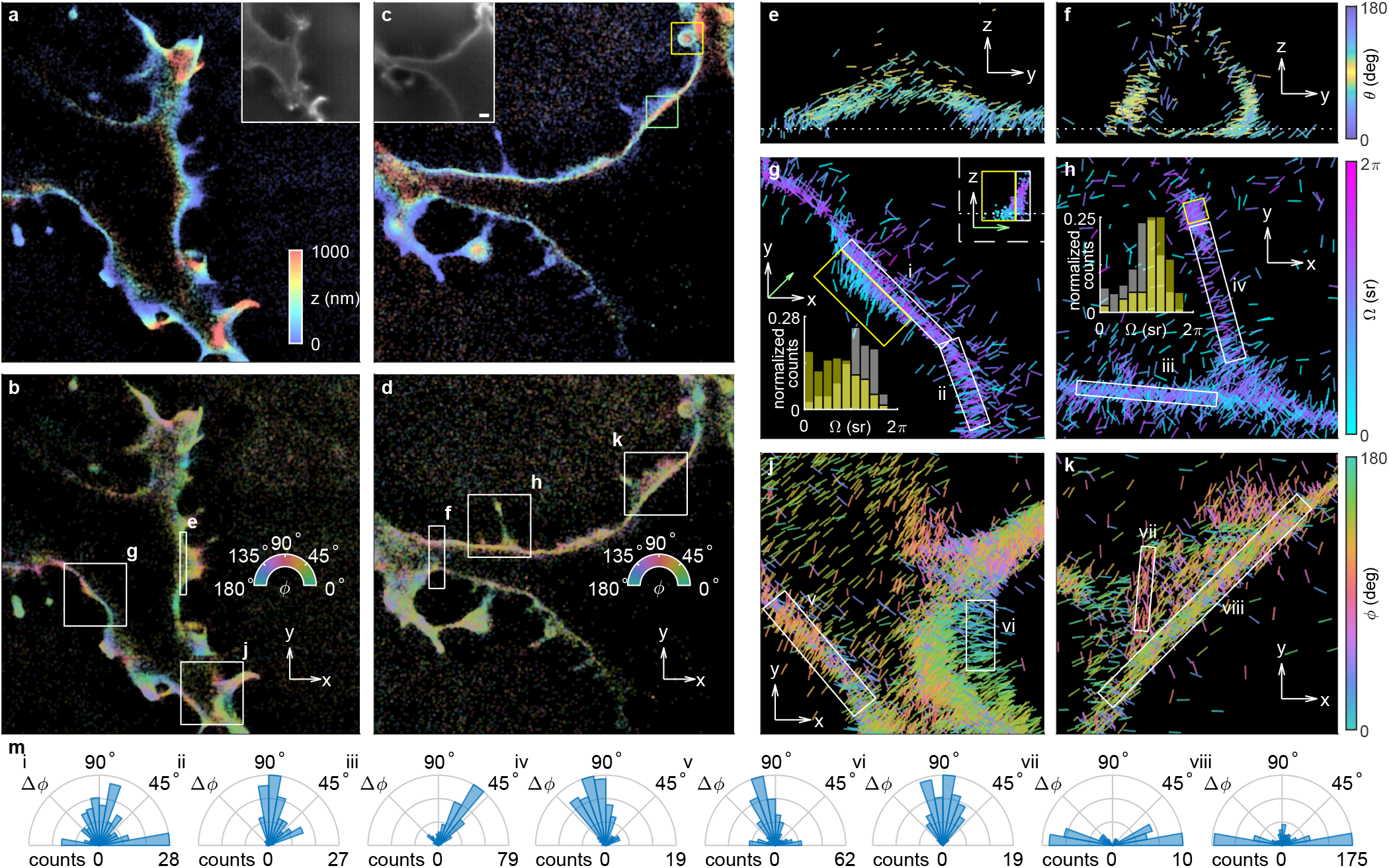
6D nanoscopic imaging MC540 molecules bound to membranes of HEK-293T cells. (a-d) Super-resolved images of two representative cells. Colors represent the estimated (a,c) axial position *z* and (b,d) azimuthal angle *ϕ*. Insets: diffractionlimited images. Scale bar: 2 µm. (e,f) *yz*-and (g-k) *xy*-views of boxed regions in (b,d). Lines are oriented and color-coded according to the measured (e,f) polar angle *θ*, (g,h) wobble angle Ω, and (j,k) azimuthal angle *ϕ*. Their lengths are proportional to (e,f) 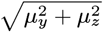 and (g-k) sin *θ*. Scale arrows: 2 µm in (a-d), 500 nm in (e-k). (g,h) Insets show the distribution of measured wobble angle Ω within the yellow and white boxed areas (i-iv). (g) Inset shows a side view of yellow and white boxed areas (i). Three-dimensional animations of the localizations in (e-k) are shown in Movie S4. (m) MC540 tilt Δ*ϕ* relative to the membrane surface within the white boxed areas in (e-h).

First, we examine the rotational mobility of MC540 molecules across various regions of the membrane. We observe smaller MC540 wobble (2.12 ± 1.40 sr, median ± std. dev., Figure 5g) near the cell-coverslip interface compared to that for membranes above the coverslip (3.62 ± 1.17 sr, Figure 5g), suggesting that the rigidity of the coverglass reduces membrane fluidity and thus constrains the rotational motions of MC540. In contrast, the tip of a membrane protrusion exhibits larger wobble angles (3.83 ± 0.98 sr, Figure 5h) compared to its middle (3.11 ± 1.16 sr, Figure 5h), implying that the smaller radius of curvature at the tip is correlated with increased intermolecular spacing and membrane fluidity. ^51^

Next, we quantify the tilt of MC540 relative to the membrane surface (within the *xy*-plane) as Δ*ϕ* = *ϕ* − *ϕ*_membrane_ ∈ [0°, 180°], where *ϕ*_membrane_ is computed using first-order LMSE fits of the localizations in regions (i)-(viii) in Figure 5g-k. Similar to NR bound to DPPC SLBs (Figure 3), cross-sectional SMOLM images (Figure 5e,f) show that MC540 binds perpendicularly to the membrane surface with tilt given by Δ*ϕ* = 101.1°± 29.9° in Figure 5h,m(iv) and Δ*ϕ* = 88.9°± 22.1° in Figure 5j,m(vi). We also detect MC540 molecules binding parallel to the membrane (Figure 5e) with a tilt of Δ*ϕ* = 0.6° ± 23.8° in Figure 5k,m(vii) (see Figure S24c,d for raw images). NR4A, ^48^ a NR variant with a membrane-targeting moiety (Figure S15), does not exhibit parallel binding behavior (Figure S25).

Interestingly, certain areas show both parallel and perpendicularly oriented MC540 [Figure 5j,m(v): 30% parallel (Δ*ϕ* = 4.6°± 18.3°) and 70% perpendicular (Δ*ϕ* = 103.2°± 18.7°) and Figure 5k,m(viii): 28% parallel (Δ*ϕ* = 83.6°± 20.8°) and 72% perpendicular (Δ*ϕ* = 0.4° ± 17.0°)]. The ratio of parallel to perpendicularly oriented MC540 even varies within a continuous region of the membrane (Figure 5g); 37% parallel (Δ*ϕ* = 6.1°± 15.1°) vs. 63% perpendicular MC540 (Δ*ϕ* = 89.7° ± 25.5°) in Figure 5g,m(i) morphs into virtually completely perpendicular MC540 in Figure 5g,m(ii) (Δ*ϕ* = 78.0°± 28.3°). Ensemble experiments have also found MC540 to be aligned both parallel and perpendicular to model membranes, ^52,53^ but because of its SM sensitivity and nanoscale resolution, raMVR imaging resolves spatial heterogeneities in the fluidity of HEK-293T membranes. The variations in MC540 orientations likely stem from ^54^ variations in lipid composition, phase, and packing of the membrane, ^55,56^ charges within the membrane, ^52^ and consequent variations in the concentration of MC540 molecules. ^57^

## Discussion

In summary, we demonstrate the design of a radially and azimuthally polarized multiview reflector (raMVR) microscope for 6D imaging of fluorescent molecules in SMOLM. When imaging Nile red binding to spherical supported lipid bilay-ers (Figure 3), we observe rotationally-fixed molecules perpendicular to the membrane and achieve a 3D experimental localization precision of ∼16 nm (std. dev., corresponding to an FWHM of 42.6 nm for 350-nm spheres) with 5000 signal photons detected. We reconstruct nearly perfectly spherical SLBs with radii from 150 nm up to 1 µm, thereby confirming the raMVR system’s robustness against refractive index mismatch. We also resolve, for the first time to our knowledge, Aβ42 aggregates intercalating with lipid membranes in 3D with nanoscale resolution (Figure 4); NR molecules bound to mixed oligomer-lipid membranes typically exhibit larger rotational motions compared to those bound to pure lipid membranes, whereas NR bound to fibrils are more fixed and parallel to the backbone. Finally, 6D SMOLM of fixed HEK-293T cells (Figure 5) resolves variations in membrane fluidity throughout the cell. Despite the relative complexity of natural cell membranes, we detect MC540 binding both parallel and perpendicular to the membrane, similar to behaviors previously observed on model membranes. ^52,53^ Notably, we also observe MC540 molecules that are neither parallel nor perpendicular to the membrane [Δ*ϕ* = 62.2° ± 31.3° in Figures 5h,m(iii)]. This observation raises the intriguing possibility that MC540 can be used to interrogate complex molecular interaction networks within biological cell membranes, such as those mediated by the cytoskeleton ^13^ and membrane proteins; ^1^ these phenomena will be investigated in future studies.

The heart of the raMVR design is the integration of a vortex waveplate and pyramidal mirrors to precisely manipulate and split polarized SM fluorescence into multiple imaging channels. One disadvantage of this design is its alignment complexity; we provide detailed schematics (Figures S1-S4) and alignment protocols (Supplementary Section 1) to mitigate these challenges. However, much like telescopes, the reflective geometry has outstanding and unique advantages for nanoscopic imaging of dim light sources; splitting both signal and background into separate channels enables 6D position and orientation measurements with minimal light loss, a large ∼2 µm depth range, and a relatively compact DSF. The raMVR mirror is specifically designed to encode molecules’ 3D positions into lateral shifts of the DSF [Figure 1e(iv,v)] and their 3D orientations into the relative brightness of the DSF in each channel (Figure S6). Thus, unlike most other methods, position and orientation information are disentangled, and the combined 6D measurement is robust across a range of *in vitro* and cellular targets. This strategy yields isotropic 3D spatial resolution, with raMVR achieving ∼26-59% smaller standard deviations on average for measuring the 3D positions and orientations compared to recent, state-of-the-art methods. ^11–13,15,17^We believe the raMVR pupil-splitting approach has many exciting possibilities for further development. It can be adapted for real-time orientation-resolved epifluorescence microscopy (Figure S14) of cellular structures. Since the 3D information within raMVR images is encoded similarly to that of SM light-field microscopy, ^58^ utilizing a fast 3D deconvolution algorithm should enable high-throughput polarization-resolved 3D imaging with larger FOVs and higher temporal resolution than demonstrated here. In addition, light-sheet illumination ^59,60^ should reduce background autofluorescence for extended-depth imaging. Finally, rapid advancements in deep learning for SM microscopy ^61–64^ could enable real-time 6D SMOLM imaging of cells and tissues, shedding light on a variety of cellular processes with unprecedented detail.

## Methods

### Imaging system

A 100x, 1.5 NA TIRF objective lens (OL, Olympus UP-LAPO100XOHR) and an achromatic tube lens (TL, *f* = 175 mm) create an intermediate image plane (IIP). Two 4f systems (L1, L2 and L3-L5, *f*_4f_ = 150 mm) then project the intermediate image planes to the cameras and create conjugate pupil planes for the waveplates and MVR mirrors of the system. A variable waveplate (VaWP, Thorlabs LCC1223-A, Figure 1a) and a vortex half waveplate (VWP, Thorlabs WPV10L-633, Figure 1a) are placed before and at the back focal plane (BFP), respectively, in the first 4f system (L1, L2). They convert radially and azimuthally polarized light to *x*-and *y*-polarized light, compensating for the birefringence of the dichroic mirrors used in our experiments ^17,30^ and are calibrated and aligned according to the procedure reported previously. ^17^ A polarizing beamsplitter (PBS, Meadowlark Optics BB-100-VIS) is placed in the second 4f system (L3-L5, Figure S4a) to create separate imaging channels containing radially or azimuthally polarized fluorescence collected from the sample. After being separated by the MVR mirrors, the polarized fluorescence is captured by detectors C1 [Hamamatsu ORCA-flash4.0v2 C11440-22CU, Figure S4a(i)] and C2 [Hamamatsu ORCA-flash4.0v3 C13440-20CU, Figure S4a(ii)]; the effective pixel size is 66.86 × 66.86 nm^2^ in object space. The detailed dimensions and placement of the pyramid mirrors are documented in Supplementary Section 1.

### Forward imaging model

For a molecule located at position ***r*** = [*x, y, z*]^⊤^ with orientational second moments ***m*** = [*m*_*xx*_, *m*_*yy*_, *m*_*zz*_, *m*_*xy*_, *m*_*xz*_, *m*_*yz*_]^⊤^, the image *I*(*u, v*) formed by a microscope is given by

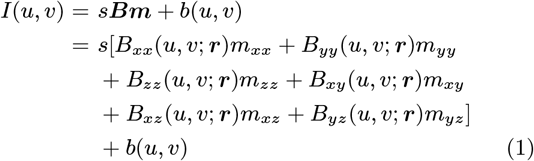

where *b*(*u, v*) represents the background fluorescence from the sample, and *B*_*ij*_ (*u, v*; ***r***) are termed the basis images (Figure S6) of the imaging system that depend on the position of the molecule (see Supplementary Section 2.1 for details). We note that the BFP segmentation guarantees that the basis images in the image plane are linearly independent with respect to the six second moments (Figure S6). The DSF in each channel translates along the *x* direction as an emitter translates in the *x* direction, as shown by the spatial gradient of each basis image [Figure S7(i-iii)]. However, if an emitter translates axially, each channel shifts laterally in a unique direction [Figure S7(iv-vi)], thereby enabling 3D positions to be estimated from the lateral positions of corresponding DSFs across all 8 channels. We also note that the direction of the *k*-vector of each channel is tilted by 1° relative to the camera normal. Thus, the apparent lateral shift caused by the non-normal propagation vector is negligible compared to that caused by the partitioning of the BFP.

### Detection and estimation algorithms for raMVR imaging

To localize SMs within MVR images for single-molecule orientation-localization microscopy (SMOLM), we need to group the DSFs detected across the 8 imaging channels to their corresponding molecules, i.e., assign spots detected in each channel to unique (*x, y, z*) positions corresponding to each emitter. Further, depending on the orientation of a particular emitter, one or more channels will contain no or very few photons, e.g., channels 6 and 8 for a molecule with orientation (*θ* = 90°, *ϕ* = 0°, Ω = 0 sr) [Figure 1b(i)] or channels 2 and 4 for a molecule with orientation (*θ* = 90°, *ϕ* = 45°, Ω = 0 sr) [Figure 1b(ii)]. While these “empty” channels improve the raMVR system’s orientation sensitivity, because these channels are only dark for specific orientations, this phenomenon makes spot assignment more challenging.

Briefly, detecting and localizing SMs follows a three-step process. First, a regularized maximum likelihood estimator is used to detect and localize SMs independently in each imaging channel (Supporting Section 3.1). Next, the coarse 3D position of each SM is calculated by grouping DSF images together across imaging channels. Finally, the 3D position and 3D orientation of each SM is estimated using maximum likelihood estimation (Supporting Section 3.2). This approach yields excellent estimation precision and accuracy (Table S2).

### Sample preparation

#### Lipid-coated silica spheres

We follow previously reported procedures ^16,17,22^ for preparing lipid-coated silica spheres. First, we prepare large unilamellar vesicles (LUVs) composed of DPPC and 40% cholesterol. A mixture of 17.62 µL of the stock DPPC solution (25 mg/mL DPPC in chloroform, Avanti Polar Lipids 850355C) and 15.46 µL of 10 mg/mL cholesterol (SigmaAldrich C8667) in chloroform are placed under vacuum overnight to evaporate the chloroform. The dried lipid mixture is then resuspended in 1 mL Tris Ca^2+^ buffer (100 mM NaCl, 3 mM Ca^2+^, 10 mM Tris, pH 7.4) to arrive at a lipid concentration of 1 mM. The lipid solution is kept under nitrogen in a water bath at 65 °C and vortexed for 20 seconds every 5 minutes for a total of 6 repetitions. The solution is then moved to an extruder set (Avanti Polar Lipids 610000) kept on a hot plate to maintain its temperature above the phase transition temperature of DPPC. A 25-passage extrusion is performed to improve the homogeneity of the size distribution of the final suspension.

We next dilute silica spheres (Bangs Laboratories SS02001, SS03001, SS04002, 2 g/mL) using Tris Ca^2+^ to reach a concentration of 40 µg/mL. It is further diluted in Tris Ca^2+^ according to Table S3. The diluted sphere solution is warmed by immersing it in a 65°C water bath and then mixed with the lipid LUV solution. The volume of each component (Table S3) ensures full coverage of SLBs on the surfaces of the spheres. ^65^ The mixture is kept in the water bath for 30 minutes and then gradually cooled to room temperature. We vortex the mixture every 5 minutes during the entire process. Finally, we centrifuge the mixture at a speed according to the sphere size (Table S3) and replace the Tris Ca^2+^ supernatant with Tris (for DPPC SLB imaging, 100 mM NaCl, 10 mM Tris, pH 7.4) or PBS (for SLBs mixed with Aβ42 imaging, 100 mM NaCl, 50 mM Na_2_PO_4_, pH 7.4).

#### Amyloid-beta 42 aggregation

The Aβ42 sample is prepared according to previously reported procedures. ^14,15^ To image the interaction between model lipid and amyloid, we incubate 20 µM monomeric Aβ42 and 1 µg/mL lipid-coated 350-nm spheres at 37 °C with 200 rpm shaking. For imaging Aβ42 fibrils on coverglass, only 10 µM monomeric Aβ42 is incubated for 48 hours.

#### HEK-293T cell culture

HEK-293T cells (ATCC CRL-3216) are maintained in Dulbecco’s modified eagle medium (DMEM, Thermo Fisher 10569044) supplemented with 10% fetal bovine serum (Thermo Fisher 26140079) and 1X Antibiotic-Antimycotic Solution (Corning 30004CI). To prepare fixed cell sample, we plate HEK-293T cells at 50% confluency on 8-chamber coverglass (Cellvis C8-1.5H-N) treated with fibronectin (SigmaAldrich FC010), according to the manufacturer’s protocol. The cells are returned to the incubator for 24 hours before they are fixed on ice with 2% paraformaldehyde in PBS for 20 minutes.

## Acknowledgements

We thank Dr. Adam Backer and Dr. Victor Acosta for helpful discussions in designing the raMVR microscope. Amyloid-beta 42 peptide was synthesized and purified by Dr. James I. Elliott (ERI Amyloid Laboratory, Oxford, CT). Research reported in this publication was supported by the National Science Foundation under grant number ECCS1653777 and by the National Institute of General Medical Sciences of the National Institutes of Health under grant number R35GM124858 to M.D.L.

## Author contributions

OZ and MDL designed and built the raMVR imaging system. ZG, YH, and MDV prepared fixed HEK293T cells. TW developed the protocol for preparing lipid-coated silica spheres. OZ performed experiments and analyzed the data.

